# Non-invasive in vivo bidirectional magnetogenetic modulation of pain circuits

**DOI:** 10.1101/2025.03.18.644041

**Authors:** Aldana M. Antoniazzi, Santiago R. Unda, Sofya Norman, Lisa E. Pomeranz, Roberta Marongiu, Sarah A. Stanley, Jeffrey M. Friedman, Michael G. Kaplitt

## Abstract

Primary nociceptors in the dorsal root ganglion (DRG) receive sensory information from discrete parts of the body and are responsible for initiating signaling events that in supraspinal regions will be interpreted as physiological or pathological pain. Genetic, pharmacologic and electric neuromodulation of nociceptor activity in freely moving non-transgenic animals has been shown to be challenging due to many factors including the immunogenicity of non-mammalian proteins, procedure invasiveness and poor temporal precision. Here, we introduce a magnetogenetic strategy that enables remote bidirectional regulation of nociceptor activity. Magnetogenetics utilizes a source of direct magnetic field (DMF) to control neuronal activity in cells that express an anti-ferritin nanobody-TRPV1 receptor fusion protein (Nb-Ft-TRPV1). In our study, AAV2retro-mediated delivery of an excitatory Nb-Ft-TRPV1 construct into the sciatic nerve of wild-type mice resulted in stable long-term transgene expression accompanied by significant reduction of mechanical withdrawal thresholds during DMF exposure, place aversion of the DMF zone and activity changes in the anterior cingulate (ACC) nucleus. Conversely, delivery of an inhibitory variant of the Nb-Ft-TRPV1 construct, engineered to gate chloride ions in response to DMF, led to reversed behavioral manifestations of mechanical allodynia and showed place preference for the DMF zone, suggestive of functional pain relief. Changes in DRG activity were confirmed by post-mortem levels, immediately following DMF exposure, of the activity-induced gene *cfos*, which increased with the excitatory construct in normal mice and decreased with the inhibitory construct in pain models Our study demonstrates that magnetogenetic channels can achieve long-term expression in the periphery without losing functionality, providing a stable gene therapy system for non-invasive, magnetic field regulation of pain-related neurons for research and potential clinical applications.

## Introduction

Pain is a complex, multifaceted experience characterized by both sensory and emotional dimensions and is often associated with actual or potential tissue damage(*1*). Various types of pain can be classified into acute, chronic, nociceptive or neuropathic, each presenting distinct challenges in clinical diagnosis and management. According to the International Association for the Study of Pain (IASP), neuropathic pain arises from a lesion or disease of the somatosensory nervous system(*1, 2*). Epidemiological studies indicate that chronic neuropathic pain affects approximately 6.9% to 10% of the general population, thereby imposing a substantial burden on global public health(*3, 4*). Despite advances in pain management, current therapeutic options frequently fail to provide adequate relief(*5*).

In recent years, breakthroughs in gene therapy have paved the way for innovative neuromodulation techniques that enable reversible, stimulus-triggered control of neuronal activity(*6–10*). Optogenetics is a popular method for circuit interrogation, with specific wavelengths of light regulating function of light-sensitive channels or signaling molecules (opsins). Optogenetic techniques necessitate the delivery of continuous light to activate opsins, a requirement that can result in local tissue heating, phototoxicity, and transient alterations in cellular phenotypes due to the use of non-endogenous proteins (*12, 13*). Although peripheral applications such as pain can use external light rather than implants used for intracranial optogenetics, the need for focal light application on the affected body part can create challenges for many assays in freely moving animals. Chemogenetics use pharmacological neuromodulation with either a small molecule ligand (Designer Receptors Exclusively Activated by Designer Drugs, DREADDs) or ionic conductance (Pharmacologically Selective Actuator Modules, PSAMs) leading to inhibition or excitation of neural activity(*11*). These can be influenced by ligand bioavailability and the inherent pharmacokinetics of the drugs used, which restrict the temporal precision and consistency of neuronal modulation. The development of a non-invasive and temporally precise neuromodulator tool could be transformative to address some of the challenges presented by these established regulatable gene therapy approaches. Magnetogenetics has recently emerged as an innovative, non-invasive alternative that addresses many of these limitations. By employing direct magnetic field (DMF) stimulation, this approach can modulate neuronal activity deep within tissue without the need for implanted hardware or continuous external stimulation(*14–18*).

Here we introduce a novel magnetogenetic technique designed to achieve precise, temporally controlled activation of pain-related neural circuits. We recently described a magnetogenetic system for regulation of specific brain circuits in freely moving mice, using an AAV expressing a nanobody (Nb) specifically directed against endogenous ferritin (Ft), fused to the TRPV1 channel (Nb-Ft-TRPV1). Expression of this activating construct in striatal D2 neurons led to freezing and neuronal activation in a magnetic field, while an inhibitory variant expressed in the subthalamic nucleus of parkinsonian mice resulted in reduced neuronal activity and resultant improvement in motor behaviors. Since this system combines domains from different native mammalian proteins, we hypothesized that this could afford advantages for manipulation of peripheral neurons, particularly those mediating pain. Using AAV2retro to deliver our excitatory Nb-Ft-TRPV1^Ca2+^ construct to pain-sensitive peripheral neurons lead to robust activation of pain circuits only when exposed to the DMF, as evidenced by behavioral evidence of pain and significant changes in *cfos* expression in both peripheral and central nervous system structures. This effect was stable to Delivery of the inhibitory Nb-Ft-TRPV1^Cl-^ construct to these same neurons in a mouse model of pain resulted in effective inhibition of neuronal activity, suppression of pain behavioral responses and reduced expression of *cfos* in dorsal root ganglia (DRG) neurons during DMF exposure. These observations highlight the dual capacity of our magnetogenetic system to both activate and inhibit pain pathways, offering a versatile tool for experimental pain research and a potential novel regulatable gene therapy for relief of intractable pain.

## Results

### Efficient delivery of transgenes to Dorsal Root Ganglia with Retrograde AAVs

Previous studies have shown successful targeting of dorsal root ganglia (DRG) neurons via sciatic injections using adeno-associated viruses (AAVs)(*19–23*). Attempting to increase efficient retrograde transport of AAV particles from axons to cell bodies in the DRG we performed intra-sciatic nerve injections of a Retrograde AAV (AAV2retro) vector encoding mCherry, under the control of the human synapsin-1 promoter (hSyn) in male C57BL/6 mice. 6 weeks post-injection, we observed robust mCherry expression in DRGs from all lumbar segments, particularly in L4 and L5 segments (L3= 3 ± 2.2, L4=29.2 ± 6.2 p=0.0009, L5=28.6 ± 9 p=0.01, L6= 15.5 ± 7.7, n.s. mCherry+ neurons per DRG) (**Figure 1a-b**). To better understand the efficiency of AAV2retro in transducing pain-related neurons, we compared tropism of this vector with cholera toxin subunit b conjugated to Alexa fluor 594 (CTb-594) and 2 mosaic hybrid serotypes (AAV2retro/rh10 and AAV2retro/6). Small-diameter neurons (<300 µm²), primarily C-fibers, are unmyelinated sensory neurons which play a crucial role in nociceptive processing. We found that out of the total number of targeted neurons, most belong to small diameter fibers, CTb-594= 78.75 ± 8.2%, AAV2retro/rh10= 79.18 ± 12.5%, AAV2retro/6= 77.5 ± 9.1%, and AAV2retro: 88.49 ± 3.0% (**Figure 1c-e**).

**Figure 1.**
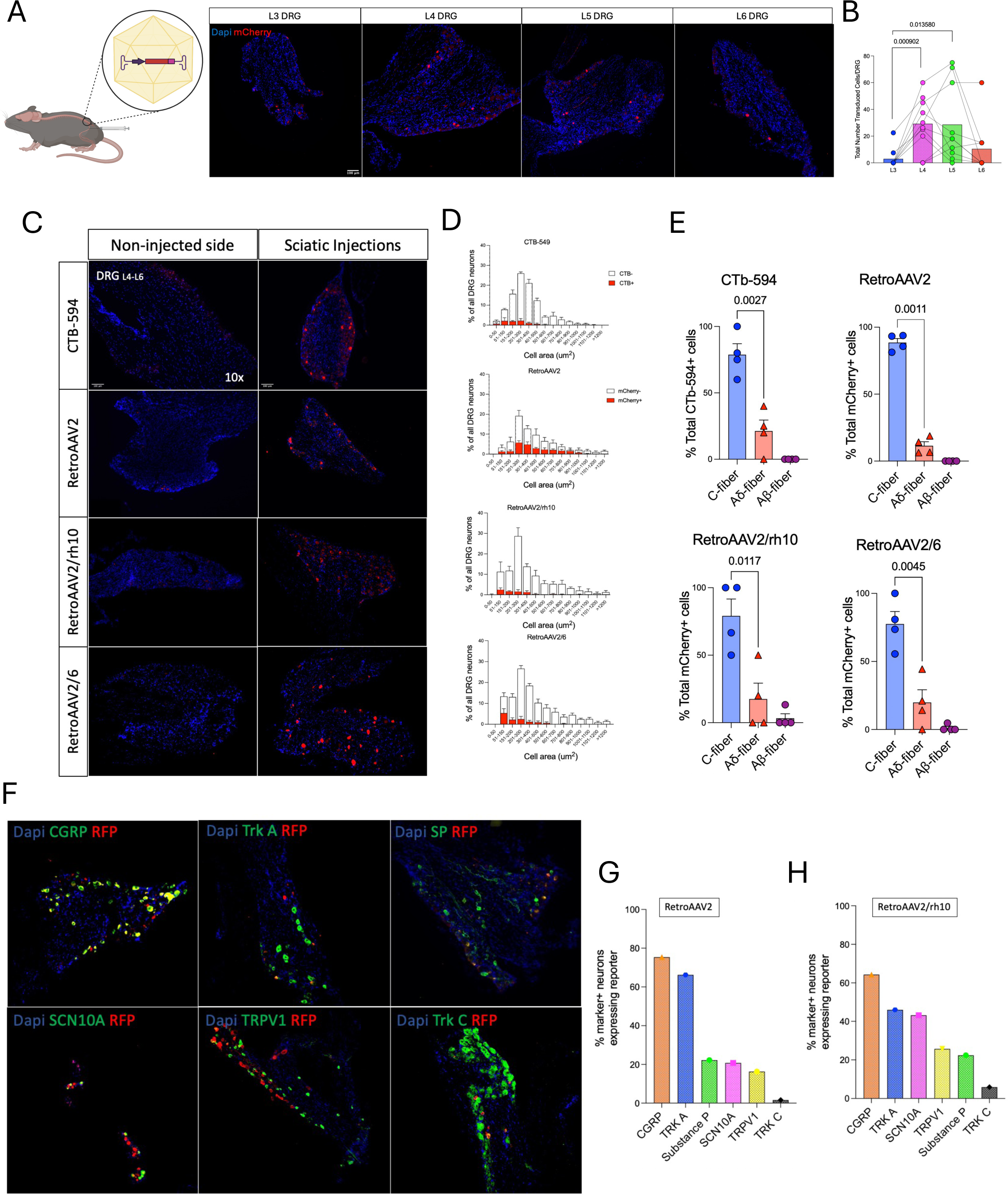
Efficient targeting of pain-related neurons by Retrograde AAV. (**A**) Anatomical target of AAV2retro in lumbar segments (L3-L6) based on total number of mCherry-expressing neurons in each DRG with quantification (**B**) (n=6-7). (**C**) Mosaic hybrid AAVretro capsid test shows mCherry transduction levels in CTb-594, AAV2retro, AAV2retro/rh10 and AAV2retro/6 injected mice. (**D-E**) Shows cell diameter histogram distribution in transduced (red bars) and non-transduced (white bars) DRG neurons and quantification of percentage of transduced cells that belonged to c-fibers, Aδ-fibers or Aβ-fibers (n=4 per capsid). (**F-G**) Representative staining of pain-related markers (in green) with transduced cells (RFP) and quantification of marker+ neurons expressing RFP in AAV2retro and AAV2retro/rh10.

Next, we assessed the co-localization of transduced cells with markers of pain-related genes in the 2 serotypes with highest efficiency in targeting small fibers (AAV2retro and AAV2retro/rh10). In AAV2retro-transduced neurons, we observed significant co-expression with calcitonin gene-related peptide (CGRP, 75.4%), tropomyosin receptor kinase A (TrkA, 66.2%), Substance P (22.17%), sodium voltage-gated channel alpha subunit 10 (SCN10A, 20.7%), and transient receptor potential vanilloid 1 (TRPV1, 16.2%), while minimal overlap was detected with tropomyosin receptor kinase C (Trk C, 1.6%). Similarly, the AAV2retro/rh10 group exhibited preferential transduction in nociceptive populations, with CGRP (64.3%), TrkA (46.03%), SCN10A (43.2%), TRPV1 (25.7%), and Substance P (22.4%), while maintaining low transduction of Trk C-positive neurons (5.9%) (**Figure 1f-h**). Then, we performed the same AAV2retro approach via hind paw injections which confirmed the AAV2retro serotype has the highest efficiency in transducing small fibers, particularly CGRP (74.9%) positive neurons (**Supplementary Figure 1a-c**).

### Magnetogenetic activation of DRG neurons induces Pain-related responses

Recently, we have shown successful neuronal activation centrally in the basal ganglia(*14*), and peripherally in the pancreas(*24*) using the excitatory magnetogenetics construct Nb-Ft-TRPV1^Ca2+^. To test whether activation of DRG neurons could elicit pain-related behaviors, we performed right intra-sciatic injections of constitutive AAV2retro-hSyn::Nb-Ft-TRPV1^Ca2+^-HA or control AAV2retro-hSyn::mCherry in wild type mice. Starting at 2 weeks post-injections mice were tested every 2 weeks for mechanical responses using Von Frey filaments while exposed to a source of direct magnetic field (DMF) of 900mT to 1.2T (**Figure 2a**). Transduction efficiency was similar between groups (mCherry= 266.8 ± 100.7 and the Nb-Ft-TRPV1^Ca2+ +^= 398.7 ± 166.2 cells per DRG, p= 0.3) (**Figure 2b-c**). While the 50% paw withdrawal threshold (PWT) did not change in the mCherry group, the Nb-Ft-TRPV1^Ca2+^-expressing mice had significantly lower 50% PWT starting at 4 weeks post-injection (50% PWT at 4wks mCherry=4 vs Nb-Ft-TRPV1^Ca2+^=1.2 ± 0.4 g, p=0.0001, 50% PWT at 6wks mCherry=4 vs Nb-Ft-TRPV1^Ca2+^=1.1 ± 0.6 g, p=0.001) (**Figure 2d**). To confirm DRG activation in a DMF-dependent manner, we analyzed the expression pattern of the activity-induced immediate-early gene, *cfos*, in mCherry- and Nb-Ft-TRPV1^Ca2+^-expressing mice exposed to a 900mT to 1.2T field. We found significant upregulation of *cfos* transcripts in the Nb-Ft-TRPV1^Ca2+^ group (mCherry=118 ± 15.9 vs Nb-Ft-TRPV1^Ca2+^=770 ± 221.6 *cfos*+ neurons per DGR, p=0.04) (**Figure 2e**).

**Figure 2.**
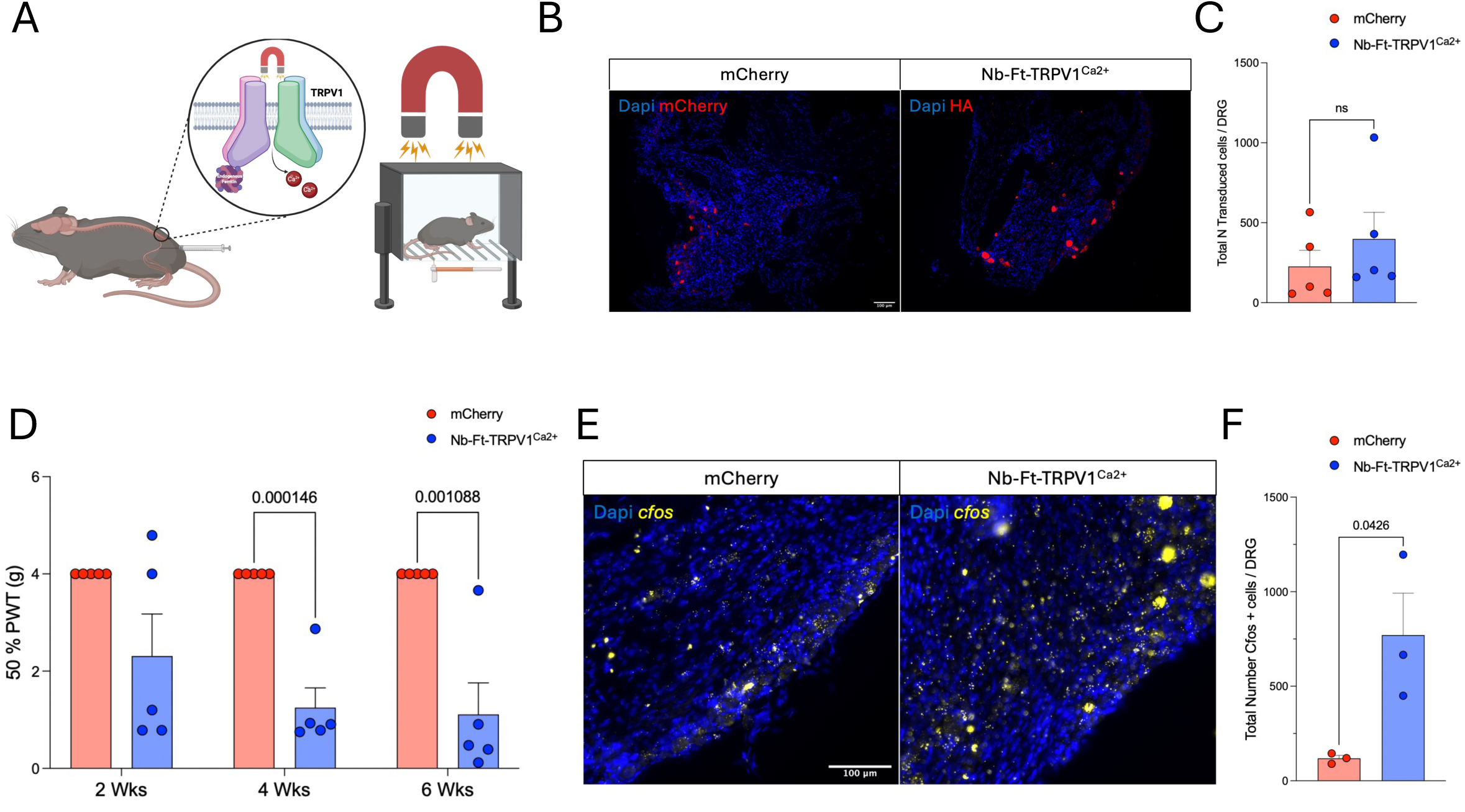
Magnetogenetic activation of DRG elicits pain-related behaviors. (**A**) Schematic figure of experimental set up. (**B**) Histological confirmation of transduced neurons in controls (mCherry, in red) and Nb-Ft-TRPV1^Ca2+^ mice (HA, in red). (**C**) Quantification of total transduced per DRG (n=5 per group). (**D**) Von Frey test at 2, 4 and 6 weeks post-intra sciatic injections of mCherry and Nb-Ft-TRPV1^Ca2+^ (n=5 per group). (**E**) Post-mortem *cfos* expression levels with DMF stimulation and quantification (**F**) in the DRG (n=3 per group). Error bars show SEM and p values were analyzed with two-tailed unpaired t-test with Welch’s correction.

In a parallel cohort of mice that received subcutaneous hind paw injections instead of intra-sciatic delivery of the transgenes, we observed similar results regarding the timing to achieve robust phenotypic changes, however overall transduction efficiency was more variable, based on the total number of transduced cells per DRG (**Supplementary Figure 1d-f**). Therefore, intra-sciatic AAV2retro delivery was used exclusively for our subsequent studies.

### Long-term stability of the excitatory magnetogenetic construct in the DRG

Previous studies have suggested optogenetics as a potential tool to modulate peripheral neural circuits(*25, 26*), however the degradation of the algae-derived opsin transgenes in the DRG after 6 to 8 weeks post-AAV delivery has been raised as a potential challenge (*13, 27*). The nanobody and the TRPV1 in our magnetogenetics constructs derive from native mammalian sequences(*14*), so we hypothesized that long-term transgene expression could be achieved in peripheral neurons mediating pain by avoiding immunogenic recognition of non-mammalian proteins. To test this, we performed side-by-side intra-sciatic injections of AAV2retro encoding the excitatory optogenetic construct (ChR2-HA) or the Nb-Ft-TRPV1^Ca2+^-HA construct at the same dose and volume. Consistent with other studies, we found that expression of optogenetic proteins almost completely lost by 8 weeks post-AAV injection, while expression of the magnetogenetic excitatory construct was maintained (ChR2= 2.5 ± 1.2 vs Nb-Ft-TRPV1= 286 ± 111, p=0.02, HA+ cells per DRG) (**Figure 3a-b**). To further test for potential immunogenicity of transduced neurons, immunostaining for nuclear factor kappa B (NF-κB) and CD45 was performed in ChR2- and Nb-Ft-TRPV1-expressing DRGs. DRG neurons expressing ChR2 had significantly increased levels of NF-κB expression compared to Nb-Ft-TRPV1-DRGs and non-injected DRGs (controls) (ChR2=1377 ± 388.5 vs Nb-Ft-TRPV1=397.3 ± 186.7, p=0.03; ChR2=1377 ± 388.5 vs control=386.7 ± 186.7, n.s. NF-κB+ cells per DRG), whereas CD45 levels were not different between ChR2 and Nb-Ft-TRPV1 groups (ChR2=140.9 ± 57.9 vs Nb-Ft-TRPV1=116.3 ± 26.4, n.s. CD45+ cells per DRG) but both constructs seemed to have increased CD45 compared to non-injected DRGs (ChR2=140.9 ± 57.9 vs control=30 ± 30 n.s; Nb-Ft-TRPV1=116.3 ± 26.4 vs control=30 ± 30, n.s. CD45+ cells per DRG) (**Supplementary Figure 2a-c**).

**Figure 3.**
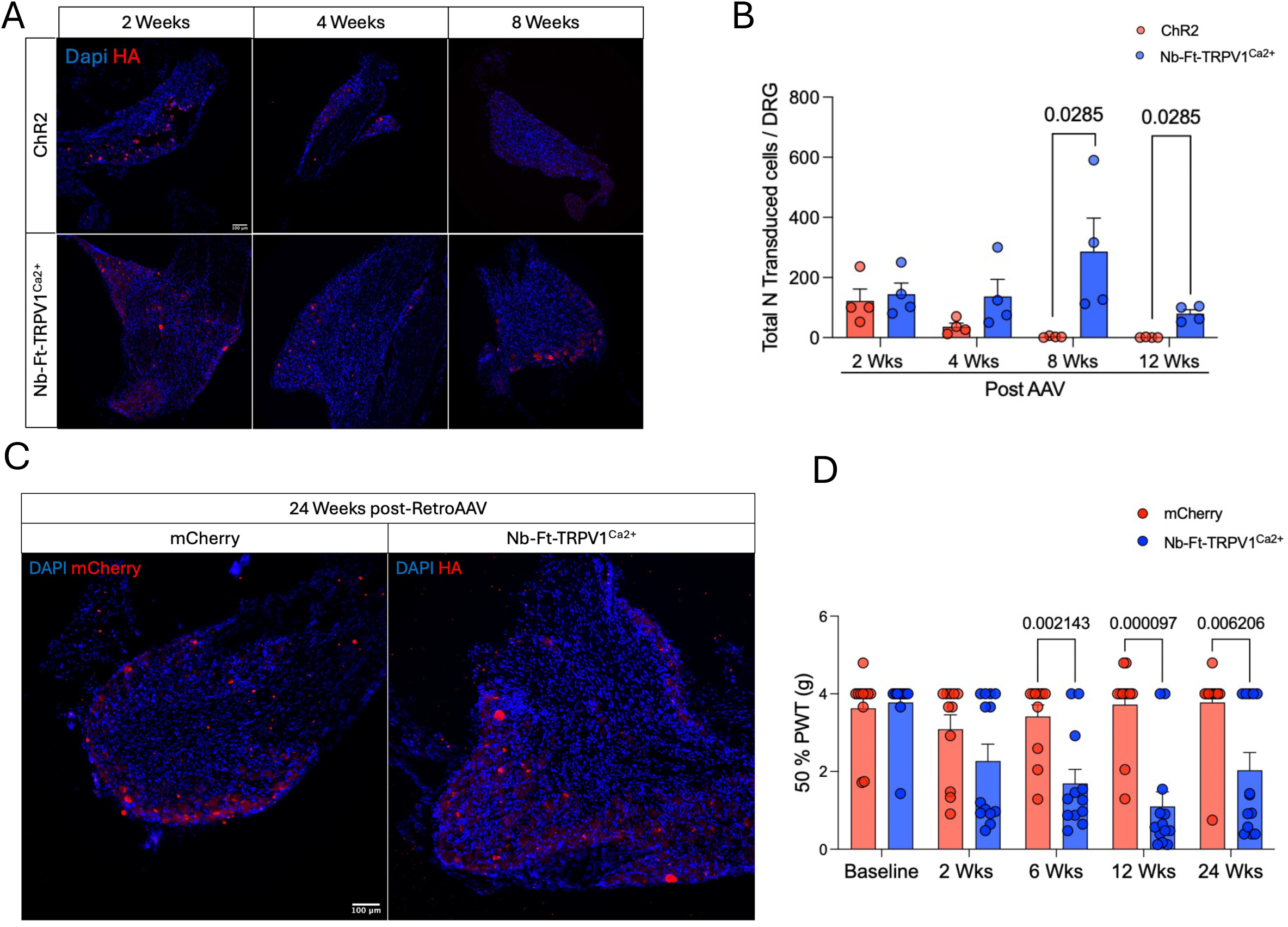
Nb-Ft-TRPV1 magnetogenetic construct achieve long-term stability and functionality in the periphery. (**A**) Histological analysis of opto- vs magneto-genetics construct overtime. (**B**) Quantification of HA+ neurons in the DRG at multiple timepoints post-intra sciatic injections (n=4 per timepoint per group). (**C**) Expression of mCherry (in red) and excitatory Nb-Ft-TRPV1^Ca2+^ (HA in red) transgenes at 24 weeks post-intra sciatic injections and (**D**) Von Frey test during DMF exposure at baseline (prior AAV2retro injection), 2, 6, 12 and 24 weeks post-AAV2retro delivery between the control (mCherry, red bars/dots) and treated (Nb-Ft-TRPV1^Ca2+^ blue bars/dots) groups (n=12 per group) expressed as 50% paw withdrawal threshold (PWT). Error bars show SEM and p values were analyzed with two-tailed unpaired t-test with Welch’s correction.

To confirm that the stable long-term expression was functionally meaningful, we tested the response of mice injected with either mCherry and Nb-Ft-TRPV1^Ca2+^-HA and exposed to a 900mT -1.2T DMF source at baseline (prior AAV2retro injections) and at 2, 6, 12 and 24 weeks post-AAV2retro delivery with the Von Frey filaments test. We observed significant differences in the 50% PWT responses in the Nb-Ft-TRPV1^Ca2+^ group starting at 6 weeks post-AAVretro and persisting to 6 months after injection (50% PWT at 6wks mCherry=3.4 ± 0.2 vs Nb-Ft-TRPV1^Ca2+^=1.6 ± 0.3 g, p=0.002, 50% PWT at 12wks mCherry=3.7 ± 0.3 vs Nb-Ft-TRPV1^Ca2+^=1.1 ± 0.3 g, p=0.00009, 50% PWT at 24wks mCherry=3.7 ± 0.3 vs Nb-Ft-TRPV1^Ca2+^=2 ± 0.4g, p=0.006). Additionally, we observed robust ongoing transgene expression histologically in the DRG at 24 weeks post-injections (**Figure 3c-d**).

### Magnetogenetic inhibition of the DRG alleviates pain-related behaviors

Recently we described a point mutation in the excitatory magnetogenetic construct that allows chloride influx instead of calcium(*14*). This functionally converted our channel to an inhibitory channel responsive to magnetic fields, which raised the potential for treatment of disorders of abnormal neuronal activation such neuropathic pain. To test the potential effect of this vector on pain responses, we performed intra-sciatic injections of AAV2retro encoding for either mCherry or the mutant Nb-Ft-TRPV1^Cl-^-HA and 6 weeks later we performed a spared nerve injury (SNI) model of neuropathic pain in the same sciatic nerve by ligating the peroneal and sural branches. Both, mCherry- and Nb-Ft-TRPV1^Cl-^-HA-expressing mice had robust transgene expression (mCherry= 97.7 ± 33.5 vs Nb-Ft-TRPV1^Cl-^-HA=84 ± 16.9, n.s., transduced neurons per DRG) (**Figure 4a-c**). Von Frey test during DMF exposure showed no differences in mechanical allodynia at baseline (prior AAV2retro) or at 4 weeks post-AAV2retro between groups, indicating that neither magnetic field alone nor the inhibitory magnetogenetic vector influenced normal responses to mechanical allodynia. However, at both 6 and 19 days post-SNI, the Nb-Ft-TRPV1^Cl-^ group showed significant alleviation of pain-responses evidenced by increased 50% PWT compared to mCherry mice (6 days post-SNI mCherry=0.4 ± 0.1 vs Nb-Ft-TRPV1^Cl-^ =2 ± 0.3 g, p=0.005; 19 days post-SNI mCherry=0.3 ± 0.06 vs Nb-Ft-TRPV1^Cl-^ =1.6 ± 0.3 g, p=0.006) (**Figure 4d**). The DRG of the Nb-Ft-TRPV1^Cl-^ group also showed a significantly decreased number of *cfos* expressing neurons compared with mCherry (mCherry=646.7 ± 133.5 vs Nb-Ft-TRPV1^Cl-^ =186.6 ± 23.1 *cfos*+ neurons per DRG, p=0.01), suggesting that exposure of the Nb-Ft-TRPV1^Cl-^ group to a magnetic field led to inhibition of DRG neuronal hyperactivity following SNI (**Figure 4e-f**). To confirm that the magnetic field was functionally regulating responses to our magnetogenetic in SNI mice, we used a place preference paradigm to determine if the zone with a sufficiently strong magnetic field was preferable. Mice were placed in a transparent long glass chamber that allows to have one zone with a magnetic field of (0.2T to 1.2T, “DMF” zone) and second zone with minimal magnetic field (<0.2T, “No DMF” zone). Recordings of mouse activity were performed at multiple time points including at baseline (prior AAV2retro) and at 3 and 6 weeks post-AAV2retro, and at 7, 15, and 23 days post-SNI. The mCherry group showed large variability in the difference between time spent in the DMF zone minus the time spent in the No DMF zone across the multiple time points prior and post SNI, indicating no evidence of preference under any condition. In contrast, the Nb-Ft-TRPV1^Cl-^-expressing mice showed clustering towards the “DMF” zone at days 7 and 15 post-SNI, consistent with the timeframe when the SNI has been described to elicit the most robust pain responses (day 7 post-SNI mCherry= -5 ± 15.03 vs Nb-Ft-TRPV1^Cl-^ =89.7 ± 17.15 sec, p=0.002; day 15 post-SNI mCherry=-39 ± 19.2 vs Nb-Ft-TRPV1^Cl-^ =35.2 ± 10.2 sec, p=0.003). At day 23 post-SNI, no differences were noted between groups, perhaps due to the known spontaneous resolution of pain in the SNI model (**Figure 4g-h**). The preference for the “DMF” zone showed a significant correlation with the transduction efficiency (Pearson’s r=0.5, two-tailed p=0.04) in the Nb-Ft-TRPV1^Cl-^ mice (**Figure 4i**).

**Figure 4.**
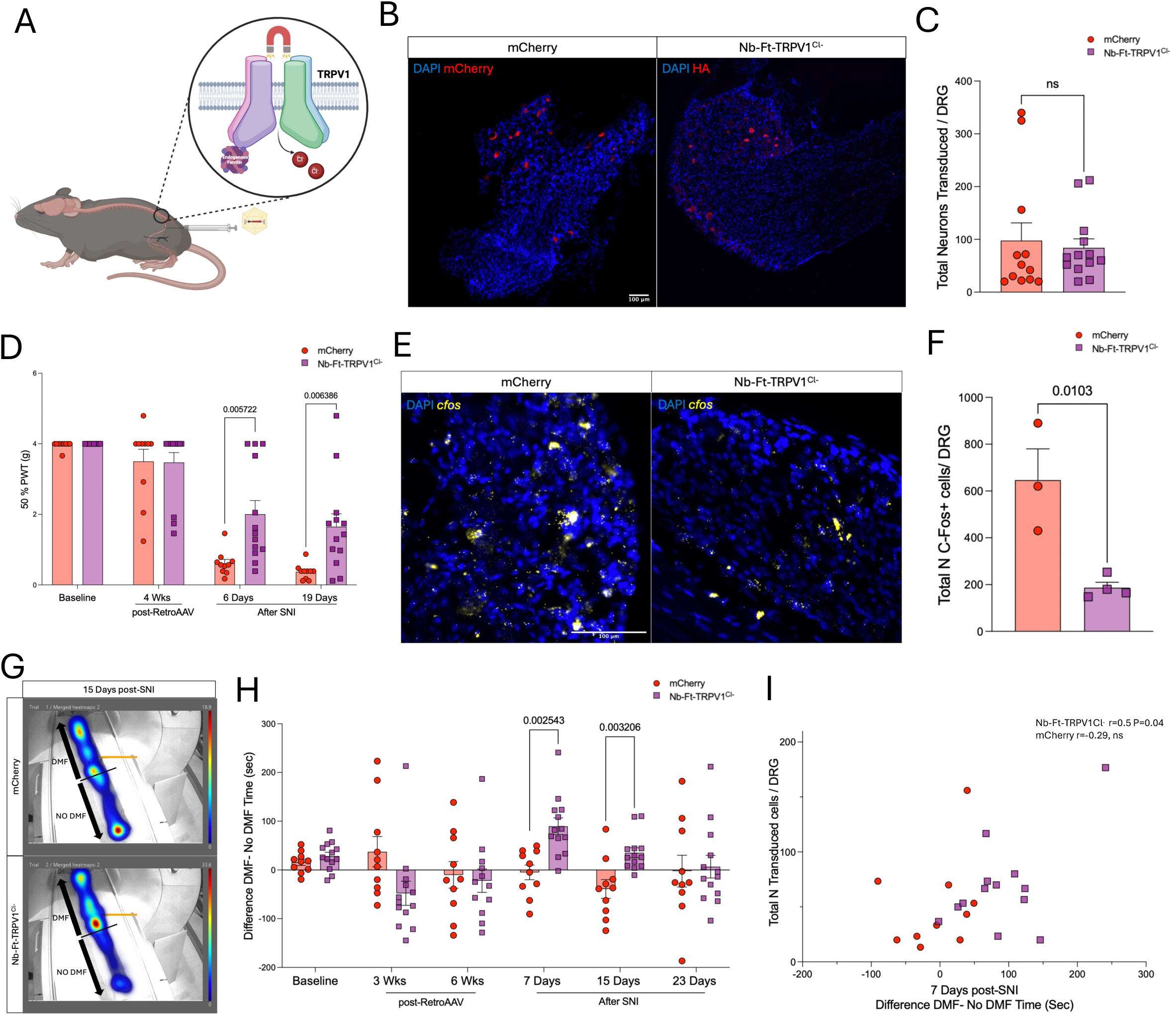
Alleviation of pain-related behaviors with the mutant inhibitory magnetogenetic construct. (**A**) Schematic figure of mutant Nb-Ft-TRPV1^Cl-^ in the DRG to gate chloride ions into the cells. (**B**) Histological confirmation of transduced neurons in controls (mCherry, in red) and Nb-Ft-TRPV1^Cl-^ mice (HA, in red). (**C**) Quantification of total transduced per DRG (n=12 per group). (**D**) Von Frey test during DMF exposure at baseline (prior AAV2retro injection), at 4 weeks post-AAV2retro delivery and at 6, and 19 days post-SNI between the control (mCherry, red bars/dots) and treated (Nb-Ft-TRPV1^Cl-^ purple bars/dots) groups (n=12/13 per group) expressed as 50% paw withdrawal threshold (PWT). (**E**) Post-mortem *cfos* expression levels with DMF stimulation and quantification (**F**) in the DRG (n=3/4 per group). Error bars show SEM and p values were analyzed with two-tailed unpaired t-test with Welch’s correction. (**G**) Heatmaps of activity in controls (mCherry) and Nb-Ft-TRPV1^Cl-^-expressing mice at 15 days post-SNI with quantification of the difference of the time spent in the DMF zone minus the time spent in the No DMF zone at baseline (prior AAV2retro), at 3 and 6 weeks (post-AAV2retro), and at 7, 15 and 23 days post-SNI (n=12/13 per group). (**H**) Correlation of total number of transduced neurons versus difference of DMF minus No DMF time. Error bars show SEM and p values were analyzed with two-tailed unpaired t-test with Welch’s correction.

In parallel, another cohort of wild type mice received intra-sciatic injections of AAV2retro encoding for the excitatory (Nb-Ft-TRPV1^Ca2+^=150.4 ± 62.1) or the inhibitory (Nb-Ft-TRPV1^Cl-^=47.7 ± 6.5, n.s. neurons transduced per DRG) magnetogenetic constructs (**Supplementary Figure 3a-c**). Transgene expression was confirmed and mechanical allodynia test with Von Frey filaments during DMF exposure showed lower 50% PWT starting at 3 weeks post-AAV2retro delivery for the Nb-Ft-TRPV1^Ca2+^ group compared to the Nb-Ft-TRPV1^Cl-^ group (50% PWT at 3wks Nb-Ft-TRPV1^Cl-^=4 vs Nb-Ft-TRPV1^Ca2+^=1.6 ± 0.5 g, p=0.002, 50% PWT at 9wks Nb-Ft-TRPV1^Cl-^=3.6 ± 0.4 vs Nb-Ft-TRPV1^Ca2+^=0.8 ± 0.1 g, p=0.0001, 50% PWT at 11wks Nb-Ft-TRPV1^Cl^ =4.2 ± 0.1 vs Nb-Ft-TRPV1^Ca2+^=0.6 ± 0.1 g, p<0.000001) (**Supplementary Figure 3d**). Analysis of *cfos* expression in these groups revealed that the excitatory magnetogenetic construct had significantly higher number of *cfos*+ cells upon DMF stimulation (Nb-Ft-TRPV1^Cl-^=267.7 ± 60 vs Nb-Ft-TRPV1^Ca2+^=892.8 ± 58.85, p<0.001 *cfos*+ neurons per DRG) (**Supplementary Figure 3e-f**). Moreover, we used the place preference paradigm to assess place aversion in the Nb-Ft-TRPV1^Ca2+^-expressing mice. The difference between DMF-No DMF spent time showed a consistent trend to avoid the DMF side of the chamber in all the time points measured in the Nb-Ft-TRPV1^Ca2+^ group, and was significant at 3 and 10 weeks post-AAV2retro (3wks Nb-Ft-TRPV1^Cl-^=91.7 ± 65.8 vs Nb-Ft-TRPV1^Ca2+^=-49.13 ± 28.3sec, p=0.05; 10wks Nb-Ft-TRPV1^Cl-^=15 ± 15.2 vs Nb-Ft-TRPV1^Ca2+^ =-59.7 ± 20.1 sec, p=0.01) (**Supplementary Figure 3g-h**).

### Magnetogenetics as a potential tool to study pain-related neural circuits

Deconstructing neural circuits associated with distinct forms of pain is crucial to understanding how brain networks are affected and what anatomical and cellular targets may be responsible for processing of specific types of pain. Since the excitatory magnetogenetic construct showed long-term stable expression and functionality in the periphery, we believe that it could be of potential interest as a tool to study acute pain responses. To explore this, we first replicated behavioral responses 6 months after intra-sciatic injections of AAV2retro encoding mCherry or Nb-Ft-TRPV1^Ca2+^ in wild type mice. There were no differences in the 50% PWT between groups at baseline (24wks post-AAV2retro without DMF mCherry=3.4 ± 0.3vs Nb-Ft-TRPV1^Ca2+^=3.4 ± 0.4 g, n.s.), but when mice were exposed to a DMF the Nb-Ft-TRPV1^Ca2+^ groups showed significant PWT reduction (mCherry=2.7 ± 0.7 vs Nb-Ft-TRPV1^Ca2+^=0.9 ± 0.1 g, p=0.01) (**Figure 5a-b**).

**Figure 5.**
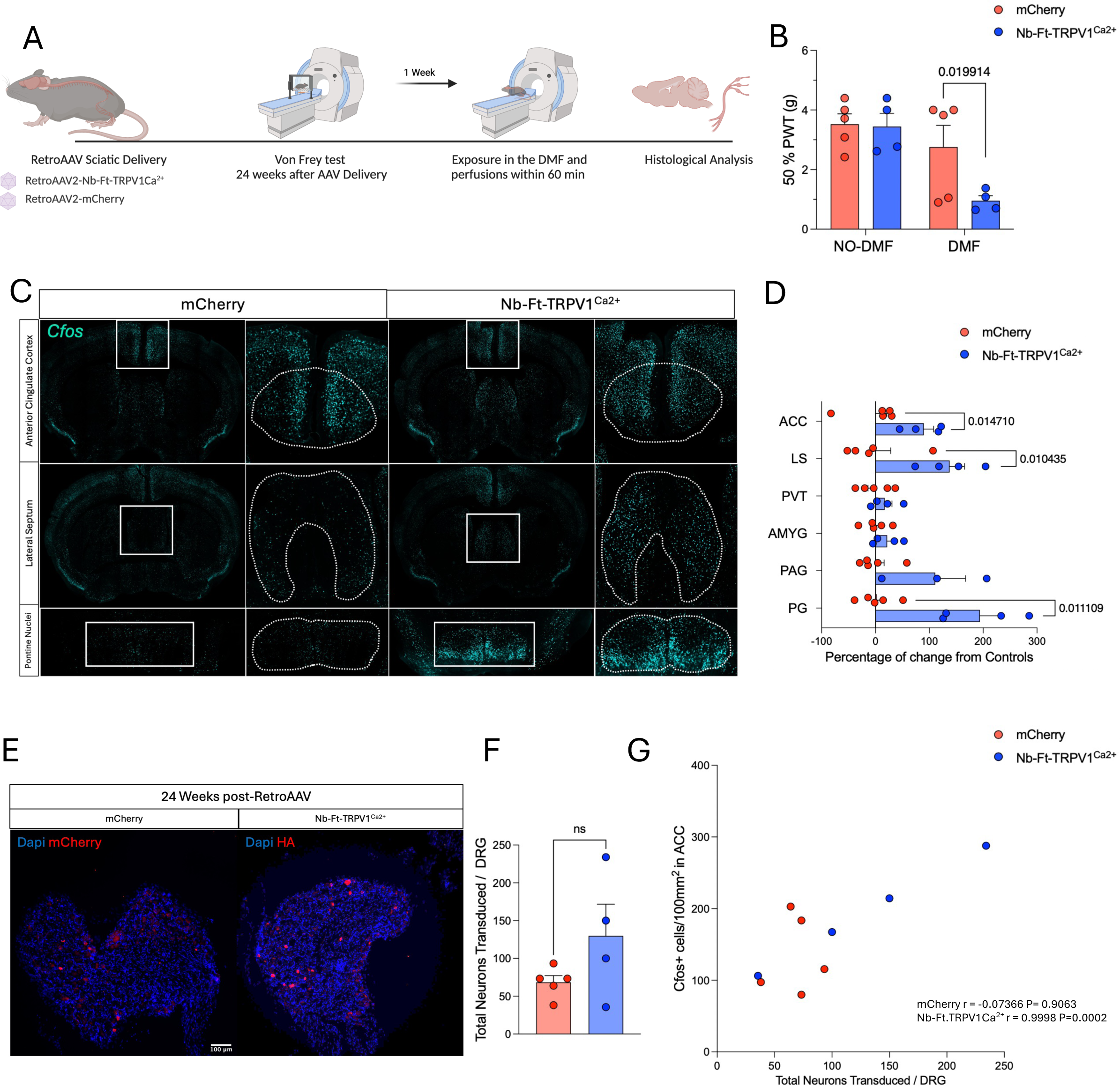
Magnetogenetic activation of DRG induces activation of pain-related brain networks. (**A**) Schematic figure of experimental set up. (**B**) Von Frey test at baseline (no DMF) and with DMF exposure in mCherry (red bars/dots) and excitatory Nb-Ft-TRPV1^Ca2+^ (blue bars/dots) expressed as 50% paw withdrawal threshold (PWT). (**C**) Representative immunostaining for *cfos* levels in the ACC, LS and PG with quantification (**D**) of *cfos* expression in ACC, LS, PVT, AMYG, PAG, and PG upon DMF exposure represented as percentage of change in the Nb-Ft-TRPV1^Ca2+^ (blue bars/dots) relative to the average expression levels in controls (mCherry, red bars/dots) (n=4/5 per group). (**E**) Histological confirmation of transgenes in the DRG with (**F**) quantification of total transduced neurons per DRG in mCherry (red bars/dots) and Nb-Ft-TRPV1^Ca2+^ (blue bars/dots). (**G**) Correlation of total number of transduced neurons versus *cfos* positive cells/100mm^2^ in the ACC. Error bars show SEM and p values were analyzed with two-tailed unpaired t-test with Welch’s correction.

After confirming robust pain induction, we analyzed the brain of these mice for activity changes upon DMF exposure. Brain regions that have been previously suggested to be part of pain-related neural circuits were examined for *cfos* expression patterns as a surrogate marker for activity, including the anterior cingulate cortex (ACC), lateral septum (LS), paraventricular nucleus of the thalamus (PVT), amygdala (AMYG), periaqueductal gray area (PAG), and the pontine gray (PG)(*28, 29*). We found that the ACC (mCherry=0.1 ± 20.9 vs Nb-Ft-TRPV1^Ca2+^=89.6 ± 18.2% of change, p=0.01), the LS (mCherry=0.1 ± 28 vs Nb-Ft-TRPV1^Ca2+^=137.7 ± 27.7 % of change, p=0.01), and the PG (mCherry=2.2 ± 15 vs Nb-Ft-TRPV1^Ca2+^=193.9 ± 39.2 % of change, p=0.01) had higher *cfos* levels the Nb-Ft-TRPV1^Ca2+^-expressing mice perhaps suggesting a role of these structures in acute pain (**Figure 5c-d**). Out of these 3 brain regions, only the ACC showed a positive correlation between the *cfos* levels and the number of transduced neurons in the DRG (Pearson’s r=0.99, two-tailed p=0.0002) while LS (Pearson’s r=0.8, two-tailed p=0.3) and PAG (Pearson’s r=0.80, two-tailed p=0.19) were not statistical different (**Figure 5e-g**). Furthermore, we proceeded to analyze the brain activity levels in SNI and Sham mice (**Supplementary Figure 4a**). First, we confirmed that the SNI mice showed robust pain responses to thermal hyperalgesia and mechanical allodynia, but no motor deficits (**Supplementary Figure 4b-d**). Then, *cfos* activity was quantified in the same brain regions as with the acute magnetogenetic activation of the Nb-Ft-TRPV1^Ca2+^ in the DRG and although there was a trend to have higher levels of *cfos* in all 6 brain structures, only the LS (SNI= 443.5 ± 208.9 vs SHAM= 0.7 ± 46.1% of change, p=0.009) and the AMYG (SNI= 216 ± 56.4 vs SHAM= 0.2 ± 36% of change p=0.02) showed significant differences compared to controls (**Supplementary Figure 4e-f**). The differences between brain activity patterns following acute Nb-Ft-TRPV1^Ca2+^ activation compared with activation patterns in the chronic SNI model supports the idea that magnetogenetics could be useful as a tool to model acute pain conditions, since it allows to for multiple trials of controlled reversible activation without inducing tissue damage and inflammation unlike other methods to induce acute pain responses(*30, 31*).

## Discussion

Recent studies have shown promising evidence that magnetogenetics can regulate central and peripheral neural circuits in a temporally and spatially precise manner(*14, 24, 32, 33*). While regulation of neural activity in animal models has been pivotal to deconstruct complex neural circuits involved in physiological and disease states, there are many challenges of tools such as optogenetics and chemogenetics including the difficulty and safety of delivering light to opsins for extended periods in animals or the poor temporal precision of small molecule ligands(*27, 34, 35*). Development of alternative strategies that allow both anatomical and temporal precision in freely moving animals could help achieve models and paradigms that better resemble the human condition, and moreover to expand the utility of these tools into suitable therapeutic applications. Here, we applied a toolkit of magnetogenetics constructs to regulate neural activity in the DRG and showed that AAV delivery of these magnetic field-based genetic sensors into the sciatic nerve are capable of inducing pain-related phenotypes upon expression of the excitatory Nb-Ft-TRPV1^Ca2+^ channel, while delivery of the Nb-Ft-TRPV1^Cl-^ inhibitory construct to peripheral neurons followed by exposure to a magnetic field resulted in therapeutic benefits in a neuropathic pain model.

Advances in AAV capsid engineering have achieved improvements in safety, targeting and performance of the vectors(*36, 37*). In recent years, in vivo directed evolution to engineer AAV capsid variants that permit retrograde uptake from projections neurons, in particular AAV2retro, have made possible to dissect circuits with greater detail(*38*). Although, axonal transport from sciatic nerve injections to DRG cell bodies has been shown with naturally occurring and modified AAVs(*20–23*), we hypothesized that the improved retrograde transport of the AAV2retro could increase transduction efficiency in transgene delivery via paw or sciatic injections. Our data showed that the AAV2retro had good transduction efficiency and tropism for small diameter fibers, mainly positive for CGRP marker, that are known to mediate nociceptive signals making this delivery approach appealing to pain research(*39*). Similar to previous studies which have elicited pain-related behaviors upon acute optogenetic activation of the DRG(*23*), acute magnetic field stimulation of the excitatory Nb-Ft-TRPV1^Ca2+^ channel using a clinically available 3T MRI device decreased the thresholds for mechanical stimuli based on the von Frey test, most likely as a result of membrane potential changes due to intracellular gating of Ca2+. Neuronal activation supported by *cfos* levels changes in DRG neurons shortly after DMF exposure. Interestingly, we found that the magnetogenetic construct has long-term stable expression and functionality in the DRG up to the latest time point analyzed (24wks post-injection). This contrasted with the almost complete loss of expression of the optogenetic construct 8wks post-injection, consistent with previous observations about potential immunogenic recognition of algae-derived Channelrhodopsin-2 (ChR2) in the periphery(*13, 23*). The mammalian nature of the TRPV1 and ferritin domains in this magnetogenetics system could represent an advantage to avoid immune-related toxicity and cell death triggered by foreign proteins as they reflect native epitopes, hence allowing for cleaner experiments that aim to understand peripheral nociceptive pathways or therapeutic approaches.

To explore the potential clinical utility of this technology, we used a mutant version of the Nb-Ft-TRPV1 construct that gates chloride into the cells. Previously we have shown that this can inhibit glutamatergic neurons in the subthalamic nucleus of parkinsonian mice resulting, in therapeutic normalization of motor abnormalities (*14*). In the current study, AAV2retro delivery of the Nb-Ft-TRPV1^Cl-^ to the sciatic nerve in the SNI model of chronic neuropathic pain resulted in higher 50% PWT in the von Frey tests when exposed to a DMF, as well as in place preference of the magnetic field zone over the non-magnetic field zone, consistent with alleviation of pain-related behaviors. Current FDA approved therapies that target the DRG to provide pain relief entail placing electrodes in close proximity to the affected DRG and a subcutaneous battery with the ultimate goal of inducing local changes in transmembrane voltage (*40*). Similar to deep brain stimulation (DBS) or spinal cord stimulation (SCS) interventions, the exact means by which therapeutic benefits of DRG stimulation (DRGS) are achieved are not fully understood. However extensive evidence that voltage-gated sodium channels (VGSC) are upregulated upon nerve injury(*41, 42*) suggests that regulation of DRG hyperexcitability may be a potential mechanism for pain relief (*43*). Hence, neuroelectric modulation of the DRG possibly exerts its effect by augmenting low-pass filtering nociceptive signals and by reducing the intrinsic excitability of neurons(*44*). This was in part demonstrated in rats with tibial nerve injury (TNI) that received DRGS, in which authors found reduction of spontaneous firing frequency of C-neurons accompanied with alleviation in spontaneous pain behaviors(*45*). Because the tropism of the AAV2retro was mainly for C-neurons, it is possible that the inhibition of small fibers and nociceptor hyperactivity is sufficient to achieve analgesic properties without interfering with the activity of myelinated fibers in the DRG. Since in our study the majority of DRG neurons transduced with AAV2retro were positive for CGRP, a well-known contributor to central sensitization(*39*), it is possible that reducing the CGRP signaling brings further biological benefits to our Nb-Ft-TRPV1^Cl-^ DRG inhibition model.

We also mapped activity changes in brain nuclei that have been described to be altered in pain conditions, both in mice with chronic neuropathic pain and in mice with acute magnetogenetic activation of the DRG. Our data showed that increased *cfos* expression in the ACC and the PG was unique to the acute magnetic field-induced activation, however only the ACC *cfos* levels correlated to the transduction efficiency, suggesting a more direct effect of the DRG activation on the ACC rather than a compensatory mechanism. This finding is supported by a recent study that described reduction in gamma-bands (30Hz) in the ACC using magnetoencephalography (MEG) to record brain activity in patients with DRGS(*46*). Furthermore, burst SCS, a type of SCS that provides pain relief by modulating the spinothalamic pathway, also affects the activity in the ACC(*47*). The spinothalamic pathway is largely modulated by C-neurons in the DRG, so the effects on the ACC are thought to be connected to activity changes in C-neurons in the DRG(*44, 48*). Our data likely reflects this peripheral to brain functional network.

In summary, we have applied a magnetogenetic toolbox to regulate peripheral pain circuits following gene delivery. Bidirectional neuromodulation of the DRG demonstrated to be useful to both, elicit and alleviate pain-related behaviors. A single AAV2retro encoding for the excitatory or the inhibitory magnetogenetics constructs targeted mainly small diameter fibers via intra-sciatic injections or paw injections which provides potential new strategies for AAV-based gene therapies with minimal invasiveness of the gene delivery to deep structures, like the DRG.

Moreover, the remote nature of the magnetic field-based gene sensors offers a paradigm of targeted neuromodulation therapy for pain without the need for a device implant. Other important findings of our work include the stability of the magnetogenetics channels in the periphery where immune cells can easily detect foreign proteins such as Channelrhodopsin. While the lack of immunogenicity is not a trivial challenge to overcome and makes the Nb-Ft-TRPV1 constructs potential candidates for future clinical applications, we acknowledge that further development of mobile devices that could create magnetic fields capable to activating magnetogenetics channels will be required to transition into suitable therapies that could alleviate pain in patients.

## Materials and Methods

### Generation of constructs

Magnetogenetics pAAV-hSyn-Nb-Ft-HA-TRPV1Ca^2+^ construct were generated as described in previous papers(*14, 24*). To generate the mutant plasmid pAAV-hSyn-Nb-Ft-HA-TRPV1^Cl-^ a 909-bp fragment encoding the I679K mutation was amplified from a TRPV1mutant construct (pSS109)(*49*) using the following primers:

Forward 5’-TCCATGGTGTTCTCCCTGGCAATGGGCTGGACCAACATGCTCT Reverse 5’-AGACTAGTGTTATTTCTCCCCTGGGACCA.

This fragment was digested with NcoI and ligated to NcoI-digested pAAV-hSyn-Nb-Ft-HA-TRPV1^Ca2+^ to generate pAAV-hSyn-Nb-Ft-TRPV1^Cl-^. The control construct pAAV-hSyn-mCherry was acquired from Addgene (catalog no. 114472). The optogenetic construct pAAV-hSyn-ChR2-HA was acquired from Vector Builder (catalog no. VB230406-1352kkn).

### Cell Culture and Adeno-Associated Virus (AAVretro) Preparation

Human embryonic kidney (HEK293, ATCC #CRL-1573) cells were cultured in DMEM (Gibco), supplemented with 10% (vol/vol) FBS (Sigma-Aldrich) and 1% (vol/vol) penicillin-streptomycin (Gibco), at 37 °C in 95% humidified air and 5% CO2. Retrograde Adeno-Associated Virus serotype 2 (AAV2retro) and its variants (AAV2retro/rh10 and AAV2retro/6) were generated via calcium phosphate-mediated transfection of HEK293 cells as previously described(*50*). Cells were harvested and lysed at 72 h after transfection. The vectors were purified using iodixanol gradient and dialyzed against PBS with 2mM MgCl2. Viral titers were quantified via quantitative real-time polymerase chain reaction (qPCR) using SYBR Green chemistry, with primers targeting the woodchuck hepatitis virus post-transcriptional regulatory element (WPRE) sequence in the AAV backbone.

### Quantitative Real Time PCR (qPCR) for Viral Titer Determination

Purified AAVretro vector preparations were processed and quantified using qPCR as described previously (1). Fast SYBR Green Master Mix (Applied Biosystems, catalog no. 4385612) was used on an AB 7500 FAST Real-Time PCR platform (Applied Biosystems), primers to the WPRE element: WPRE-Fw: 5’-GGCTGTTGGGCACTGACAAT-3’; WPRE-Rev: 5’-CTTCTGCTACGTCCCTTCGG-3’. The relative number of full viral particles were calculated using the standard curve method by normalizing to known standard samples.

### Animals and Housing Conditions

8-10 weeks old male wild type C57BL/6J mice, obtained from Jackson laboratory (Jax.org; strain #:000664) were housed five per cage and kept at 22°C on a reverse 11 am-light/11 pm-dark cycle, with ad libitum access to standard chow and water. All experimental procedures were approved by the Institutional Animal Care and Use Committee (IACUC) of Weill Cornell Medicine and complied with National Institutes of Health (NIH) guidelines (IACUC #2009-0026).

### Sciatic Nerve Injections

Surgical procedures were conducted under anesthesia using an intraperitoneal injection of ketamine (110 mg/kg, Butler Animal Health Supply) and xylazine (10 mg/kg, Lloyd Laboratories). The surgical site was sterilized with alternating applications of povidone-iodine (Betadine®) or chlorhexidine (Nolvasan®) scrub and 70% isopropyl alcohol or ethanol-soaked gauze sponges.

A posterior skin incision was made in the mid-thigh region, and the sciatic nerve was exposed via blunt dissection between the gluteus superficialis and biceps femoris muscles. Minimizing nerve manipulation, a total of 1 × 10¹ viral genomes of AAVretro suspension was injected at a rate of 1 µL/min using a 10-µL Hamilton syringe equipped with a 36G NanoFil beveled needle (World Precision Instruments). Injections were administered at two sites within the peroneal and tibial branches of the sciatic nerve to ensure efficient viral delivery. The needle remained in place for 5 seconds post-injection to prevent viral reflux. Incisions were closed using 4-0 absorbable sutures (Demetech Demecryl), and mice were allowed to recover for six weeks before undergoing behavioral assessments or Spared Nerve Injury (SNI) surgery, ensuring optimal transgene expression.

### Paw injections

Mice were anesthetized intraperitoneally with ketamine (110 mg/kg) and xylazine (10 mg/kg) and placed on a pre-warmed heating pad to maintain body temperature. The right hind paw was then cleaned with a cotton swab soaked in 70% ethanol and allowed to air dry. A sterile syringe equipped with a 30-gauge needle was used to administer the AAVretro viral vectors (titer: 5 × 10¹ viral genomes) into the plantar surface of the paw intradermally. The injection was delivered slowly over a period of 15–30 seconds to minimize tissue trauma and ensure uniform distribution of the compound. Following administration, the needle was carefully withdrawn, and gentle pressure was applied to the injection site with a sterile cotton swab to prevent leakage and control minor bleeding. After the injection, animals were monitored continuously until they fully recovered from anesthesia. Once fully alert, animals were returned to their home cages and observed for at least 3 days for any signs of distress or adverse reactions.

### Direct Magnetic field Exposure

A direct magnetic field (DMF) was generated using a Siemens 3.0 Tesla PRISMA MRI Scanner (Siemens Healthcare). The magnetic field strength was measured and mapped using a F.W. Bell Model 5080 Gaussmeter.

### Spared Nerve Injury

After 6 weeks post AAV viral delivery we performed a Spared nerve injury as a chronic neuropathic pain model. The animals were anesthetized with ketamine/xylazine at concentrations of 110 and 10 mg/kg body weight i.p., respectively. After the skin was shaved in the right thigh sterilization of the area was done with topical application of povidone-iodine prep pad and 70% isopropyl alcohol. We then made an incision at the mid-thigh using the femur as a landmark, then a blunt dissection through the biceps femoris muscle. The sciatic nerve and its three brunches were exposed. In this surgical procedure the common peroneal and tibial nerves are ligated using a 6-0 plain chromic gut and axotomize. Keep the sural nerve intact. In the sham operation, the skin and muscles were opened and immediately closed to avoid damage or manipulation of sciatic nerve. The incision was closed using 4-0 demetech demecryl sutures.

## Behavioral Assessments

### Mechanical Allodynia (Von Frey Test)

Mechanical hypersensitivity was evaluated using Von Frey filaments (Bioseb, Ref: BIO-VF-M) and the Up-Down method, as previously described(*51*). Mice were acclimated in mesh-bottom chambers for 30–45 minutes before testing. Calibrated filaments of varying forces were applied to the ventral surface of the hind paw, and withdrawal responses (paw withdrawal, toe spreading, or licking) were recorded. Threshold values were determined using Up Down Reader software(*52*).

### Place preference/aversion

To evaluate magnetically induced aversion or preference, we employed a modified conditioned place preference (CPP) assay. The testing apparatus consisted of a 1-meter-long chamber divided into two distinct zones: (1) the direct magnetic field (DMF) zone, positioned close to the 3.0 Tesla MRI magnetic field (field=1.2T-0.2T), and (2) the No-DMF zone, located outside the magnetic field (field<0.2T).

Mice were habituated to the chamber for three consecutive days (10 minutes per day) before baseline testing. On the test day, animals were allowed 2 minutes of unrestricted exploration before initiating a 5-minute recorded session to assess place preference or aversion. Behavioral assessments were conducted at 3, 6, and 10 weeks post-AAVretro delivery, as well as at 7, 15, and 23 days post-Spared Nerve Injury (SNI) to evaluate the long-term impact of viral transduction and nerve injury on magnetically induced behaviors.

Behavioral tracking was performed using Noldus EthoVision XT14 software, employing automated deep-learning-based animal detection to precisely track the center point of each mouse in real-time. The time spent in each zone (DMF vs. No-DMF) was extracted from video recordings, and heatmaps were generated using EthoVision analysis tools to visualize spatial preferences.

### Open-Field Arena Behavioral Assessment

Locomotor activity was evaluated in a standardized open-field arena to assess motor dysfunction in mice following SNI. Four identical arenas were simultaneously used for testing, and recordings were conducted at 3, 7, and 14 days post-SNI. To minimize the potential confounding effects of a novel environment, each mouse was habituated to the open-field arena for 10 minutes on two consecutive days prior to testing. On the test day, each mouse was placed in the center of the arena, and spontaneous locomotor behavior was recorded continuously for 30 minutes. Video recordings were subsequently analyzed using EthoVision XT software to quantify locomotor parameters such as total distance traveled in predefined zones.

### Immunohistochemistry of Dorsal Root Ganglia (DRG)

Following completion of all assessments, mice were deeply anesthetized with sodium pentobarbital (150 mg/kg, intraperitoneal i.p.) and transcardially perfused with 4% paraformaldehyde (PFA). The brains, spinal cords, and dorsal root ganglia (DRG) were harvested, post-fixed overnight in 4% PFA, and cryoprotected in 30% sucrose. DRGs were cryosectioned at 20 µm thickness onto Superfrost slides. To visualize viral expression and pain-related markers, sections were washed twice with Dulbecco’s phosphate-buffered saline (DPBS) and then incubated in a blocking solution containing 10% donkey serum in Tween-20 (TBS-T; 50 mM Tris, 150 mM NaCl, 0.1% Tween-20, pH 7.4) for 1 hour at room temperature. The following primary antibodies were applied and incubated overnight at 4°C in a humidified chamber: anti-transient receptor potential vanilloid 1 (Alomone Labs, catalog no. AGP-118; dilution 1:100), anti-hemagglutinin (Sigma Aldrich, catalog no. 11867423001; dilution 1:100), anti-red fluorescent protein (Rockland, catalog no. 200-101-379S, dilution 1:500), anti-tropomyosin receptor kinase A (Sigma-Aldrich, catalog no. SAB4502031-100UG; dilution 1:100), anti-tropomyosin receptor kinase C (Thermo Fisher, catalog no. 701985; dilution 1:100), and anti-substance P (Fisher Scientific, catalog no. 556312; dilution 1:400). The following day, sections were rinsed twice for 15 minutes with Tris-buffered saline containing Tween-20 (TBS-T) and incubated with species-specific Alexa Fluor-conjugated donkey secondary antibodies using a dilution of 1:500 for 2 hours at room temperature. The secondary antibodies used included donkey anti-rat IgG (H+L), Alexa Fluor 488 (Invitrogen, catalog no. A-21208); donkey anti-rat IgG (H+L), Alexa Fluor 594 (Invitrogen, catalog no. A-21209); donkey anti-goat IgG (H+L), Alexa Fluor 594 (Invitrogen, catalog no. A-11058); donkey anti-rabbit IgG (H+L), Alexa Fluor 488 (Invitrogen, catalog no. A-21206); and donkey anti-guinea pig IgG (H+L), Alexa Fluor 647 (Jackson ImmunoResearch, catalog no. 706-605-148). To minimize cross-reactivity, all secondary antibodies were highly cross-adsorbed against immunoglobulins from multiple species. After two additional DPBS washes, sections were counterstained with 4′,6-diamidino-2-phenylindole (DAPI; 1:10,000) for 10 minutes, air-dried, and cover slipped with Prolong mounting medium. For *cfos* analyses, mice were exposed to a direct magnetic field (DMF; ∼1.2 Tesla from an MRI magnet) for 20 minutes prior to perfusion, with all animals processed within one-hour post-DMF exposure.

### Immunohistochemistry of Brain Sections

Brain tissue was sectioned at 30 µm using a vibratome (Leica Microsystems). Free-floating sections were washed twice for 15 minutes with Tris-buffered saline containing Tween-20 (TBS-T; 50 mM Tris, 150 mM NaCl, 0.1% Tween-20, pH 7.4) and blocked for 1 hour at room temperature in 10% normal serum (in TBS-T). Sections were then incubated overnight at 4°C with a rat anti-*cfos* primary antibody (Synaptic Systems, catalog no. 226-017; dilution 1:500). The following day, tissues were washed twice for 15 minutes with TBS-T and incubated with an Alexa Fluor 594-conjugated donkey anti-rat IgG (H+L) secondary antibody (Life Technologies, catalog no. A-21209; dilution 1:1000) for 1 hour at room temperature. Subsequent to two additional 15-minute TBS-T washes, sections were counterstained with DAPI (1:10,000; AAT Bioquest, CAS 28718-90-3) for 5 minutes, followed by two final washes in TBS-T prior to mounting on slides and cover slipping with Mowiol. All incubations and washes were performed on a shaker.

### RNAscope In Situ Hybridization

RNAscope in situ hybridization was conducted according to the manufacturer’s protocol (Advanced Cell Diagnostics, ACDbio), with modifications to omit baking steps at 60°C. All incubations at 40°C were performed in a HybEZ oven using the EZ-Batch system (ACDbio). Sections were air-dried for 1 hour, post-fixed in 4% PFA for 15 minutes at 4°C, and then dehydrated. Tissues were treated with hydrogen peroxide (ACDbio, catalog no. 322335) at room temperature for 10 minutes, followed by incubation in target retrieval buffer (ACDbio, catalog no. 322000) for 10 minutes at 100°C. After two washes with double-deionized water (ddH O), sections were incubated in 100% ethanol for 3 minutes and dried at 60°C for 10 minutes. Protease treatment was performed using Protease III (ACDbio, catalog no. 322337) at 40°C for 30 minutes, followed by five washes with ddH O. Sections were then incubated for 2 hours at 40°C with probes targeting Mm-Calca (Channel 1), SCN10A (Channel 1), and mCherry (Channel 4). Positive (ACDbio, catalog no. 321811) and negative (ACDbio, catalog no. 321831) control probes were included on separate slides. After two washes with ACDbio wash buffer (catalog no. 310091) at 40°C, the sections underwent sequential amplification: 30 minutes with AMP1, 30 minutes with AMP2, and 15 minutes with AMP3 (ACDbio kit, catalog no. 323110). Subsequently, tissues were incubated with horseradish peroxidase (HRP) for Channel 1 (HRP C1) for 15 minutes, followed by signal detection using Opal 520 (1:1500; catalog no. OP-001001) in TSA buffer (ACDbio, catalog no. 322809). After a 15-minute HRP blocking step, a similar procedure was performed using HRP Channel 4 (HRP C4) combined with Opal 570 (1:1500; catalog no. OP-001005). Finally, slides were cover slipped using Prolong Gold Antifade Mounting solution with DAPI (ACDbio).

### Imaging Acquisition and Analysis

Fluorescence imaging was performed on an epifluorescence microscope (Olympus BX61) equipped with an Olympus DP71 digital camera. For quantitative analyses, three representative slices were randomly selected from a series of 20 cryostat sections; the average cell count per slice was extrapolated to estimate the total across the series. DRG images were acquired using a 10× objective lens, with *cfos*-positive neurons identified only upon clear co-localization with DAPI. Quantitative analyses for brain sections were conducted using ImageJ and CellProfiler software.

### Statistics

Exact sample sizes for each experiment are indicated in figure legends. The power was set at 95% probability, effect size to 0.3 to 0.5 depending on the experimental outcome, and significance at P < 0.05.

Statistical analyses were conducted using GraphPad Prism 7.0. The statistical test used in each experiment depended on the type of data collected (reported in figure legends). We tested for normality using the Kolmogorov-Smirnov test and Q-Q plots. Paired or unpaired T-test, Two-way ANOVA or two-tailed t test for statistical comparisons were performed as appropriate.

Pearson’s correlation coefficients were used to quantify associations between different readouts (e.g., transduced neurons versus behavioral response). When comparing two sets of normally distributed data, two-tailed t tests were used.

## Acknowledgments

**Funding:** This work was supported by the National Institutes of Health R01 NS097184 (J.M.F., M.G.K., and S.A.S.), National Institutes of Health OT2OD024912 (S.A.S.), and JPB Foundation (J.M.F. and M.G.K.).

**Competing interests:** S.A.S. and J.M.F. are inventors listed on patents US10786570B2 and US20210052909A1 assigned to The Rockefeller University. The other authors declare that they have no competing interests.

