## Supplementary figures for "Non-invasive in vivo bidirectional magnetogenetic modulation of pain circuits"

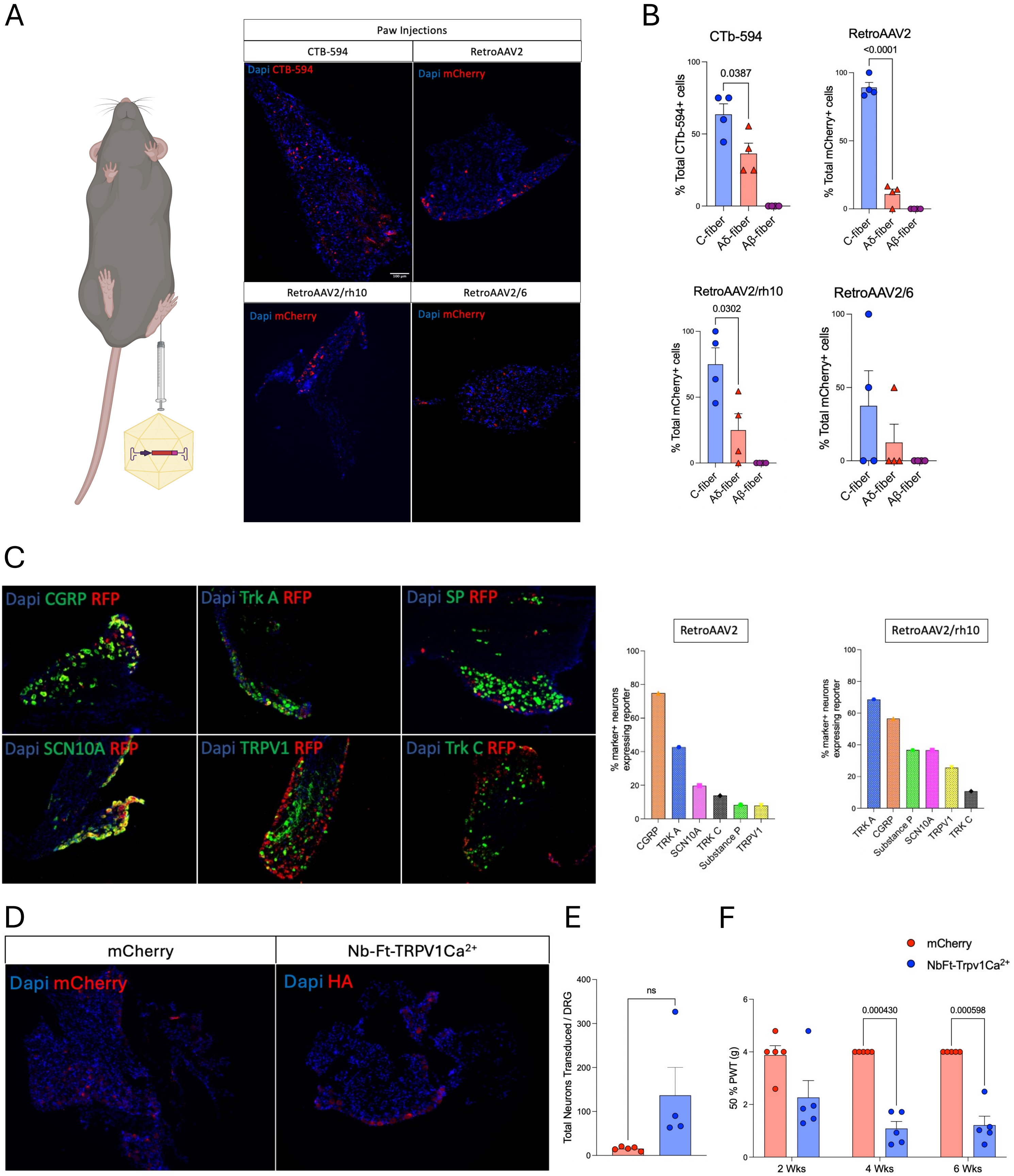


Supplementary Fig. 1

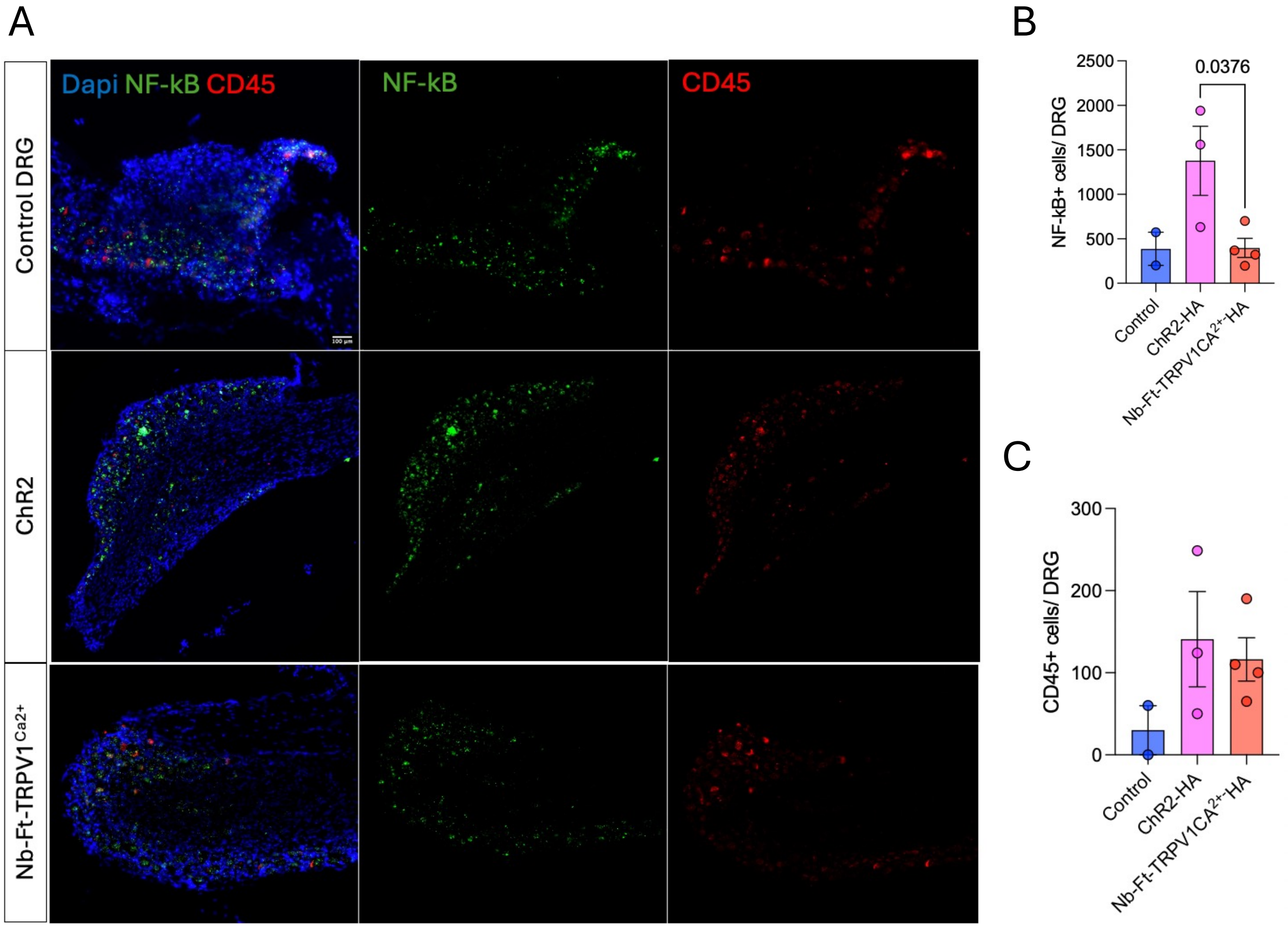


Supplementary Fig. 2

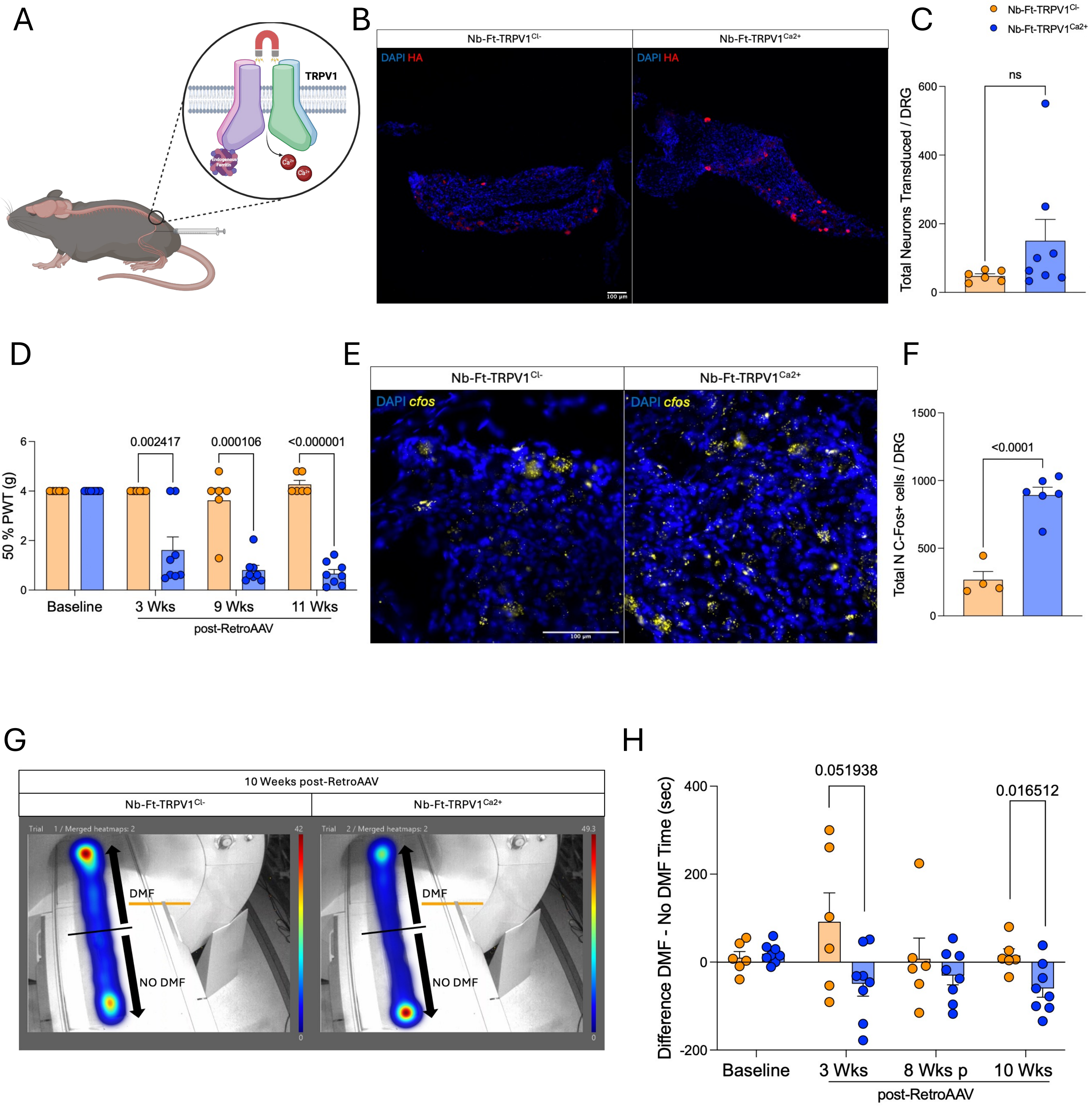


Supplementary Fig. 3

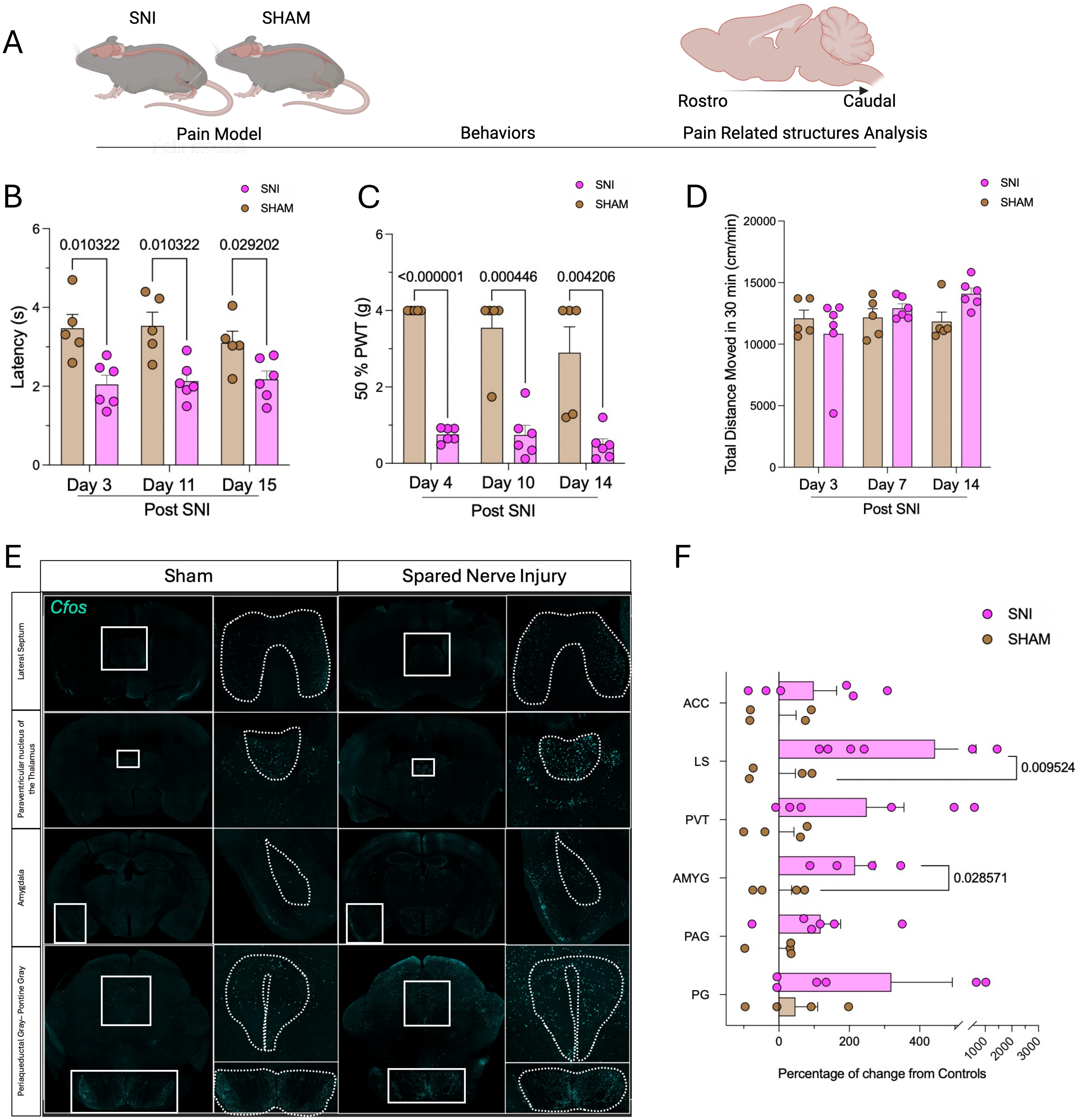


Supplementary Fig. 4

**Supplementary Figure 1. Targeted Transduction of Dorsal Root Ganglia (DRG) Neurons by RetroAAV.**

**(A)** DRG transduction following paw injection of CTb-594, RetroAAV2, RetroAAV2/rh10, and RetroAAV2/6. **(B)** Quantification of transduced neurons categorized into C-fibers, Aδ-fibers, and Aβ-fibers (n = 4 per capsid). **(C)** Representative immunofluorescence images showing colocalization of pain-related markers (green) with mCherry-expressing neurons in RetroAAV2- and RetroAAV2/rh10-injected mice. **(D)** Histological validation of transduced neurons in control (mCherry, red) and Nb-Ft-TRPV1Ca2+ mice (HA, red). **(E)** Quantification of total transduced neurons per DRG (n = 4–5 per group). **(F)** Von Frey test assessing mechanical allodynia at 2, 4, and 6 weeks post-paw injection of mCherry or Nb-Ft-TRPV1Ca2+ (n = 5 per group). Data are presented as mean ± SEM. Statistical significance was determined using a two-tailed unpaired t-test with Welch's correction.

**Supplementary Figure 2. Immunohistochemical Analysis of Inflammatory Markers in DRGs from Optogenetics, Magnetogenetics, and Control Groups.**

**(A**) Representative histological sections showing NF-κB staining from Optogenetic (ChR2), Magnetogenetic (Nb-Ft-TRPV1$C^{a2+}$), and control DRG tissues, with quantification of NF-κB–positive cells (n = 2–4 per group) **(B)**. **(C)** Quantitative analysis of CD45 expression in DRGs across the same experimental conditions (n = 2–4 per group). Data are expressed as mean ± SEM. Statistical significance was determined using two-tailed unpaired t-tests with Welch's correction.

**Supplementary Figure 3. Characterization of Nb-Ft-TRPV1 Magnetogenetic Constructs in DRG Neurons.**

(**A**) Schematic illustration of the Nb-Ft-TRPV1Ca2+ construct, designed to modulate neuronal activity via calcium gating. **(B)** Representative histological images confirming neuronal transduction in DRGs from control mice expressing Nb-Ft-TRPV1Cl– and treated mice expressing Nb-Ft-TRPV1Ca2+ (HA immunolabeling in red). **(C)** Quantification of total transduced neurons per DRG (n = 6–8 per group). **(D)** Mechanical allodynia measured by the Von Frey test under DMF exposure, at baseline (prior to RetroAAV2 injection) and at 3, 9, and 11 weeks post-injection, expressed as the 50% paw withdrawal threshold (PWT). Post-mortem *cfos* expression **(E)** and quantification of *cfos* levels in DRGs following DMF stimulation (n = 4–6 per group) **(F)**. **(G)** Heatmaps of activity at 10 weeks post RetroAAV and **(H)** analysis depicting the difference in time spent in the DMF versus the no-DMF zone at baseline and at 3, 8, and 10 weeks post-RetroAAV2 delivery, comparing control (Nb-Ft-TRPV1Cl–expressing) and Nb-Ft-TRPV1Ca2+ mice. Data are presented as mean ± SEM. Statistical significance was determined using two-tailed unpaired t-tests with Welch's correction.

**Supplementary Figure 4. Experimental Timeline and Behavioral Assessments Following Spared Nerve Injury (SNI).**

**(A)** Schematic representation of the experimental timeline. **(B)** Hargreaves test performed at 3, 11, and 15 days post-SNI in sham (brown bars) and SNI (pink bars) groups (n = 5–6) to assess thermal nociception. **(C)** Von Frey test conducted at 4, 10, and 14 days post-SNI, with results expressed as the 50% paw withdrawal threshold (PWT). **(D)** Open field test performed at 3, 7, and 24 days post-SNI (n = 5–6), with locomotor activity quantified as the total distance traveled in 30 minutes. **(E)** Representative immunostaining for cfos in the lateral septum (LS), paraventricular thalamus (PVT), amygdala (AMYG), periaqueductal gray (PAG), and parabrachial nucleus (PG). **(F)** Quantification of cfos expression in the anterior cingulate cortex (ACC), LS, PVT, AMYG, PAG, and PG is shown as the percentage change in SNI (pink bars/dots) relative to sham controls (brown bars/dots) (n = 4–5 per group). Data are presented as mean ± SEM. Statistical analyses were performed using two-tailed unpaired t-tests with Welch's correction.
